# Identification of osteopontin as a positional and functional candidate gene for cardiac hypertrophy in the SHRSP rat

**DOI:** 10.64898/2026.08.24.746886

**Authors:** Cara Trivett, Tamara P. Martin, Amrita Asirvatham, Kirsty Foote, Aiste Monkeviciute, Wendy Beattie, Christopher M. Loughrey, John D. McClure, Anna F. Dominiczak, Delyth Graham, Martin W. McBride

## Abstract

Left ventricular hypertrophy, common in cardiometabolic and renal disease, is a major risk factor for cardiovascular morbidity and mortality. Left ventricular mass is a highly heritable, polygenic trait. Linkage studies in WKY and SHRSP rats have identified a quantitative trait locus for left ventricular mass index on chromosome 14. Congenic strains, where trait-associated genetic loci are introduced into a control strain, can identify causal genetic mediators relevant to human disease.

Chromosome 14 congenic (WKY.SPGla14a), WKY, and SHRSP strains underwent cardiac phenotyping and transcriptome profiling at; 1-3 days (neonate), 5 weeks, and 16-weeks. Compared to WKY, LVMI was significantly increased in SHRSP and WKY.SPGla14a at 5 weeks (LVMI_SHRSP-WKY_=0.26g/kg, LVMI_WKY.SPGla14a-WKY_=0.30g/kg), prior to measured hypertension in this model. SHRSP blood pressure was significantly greater than WKY.SPGla14a, and WKY from 12-20 weeks (AUC_diff_=497 *vs* WKY, AUC_diff_=412 *vs* WKY.SPGla14a). Cardiac transcriptome analysis of neonate, 5-week, and 16-week hearts identified significantly increased expression of secreted phosphoprotein 1 (*Spp1/*osteopontin) in SHRSP and WKY.SPGla14a compared to WKY, which is positioned within the transferred congenic region. Overexpression of *Spp1* mRNA significantly increased H9c2 cell size and was shown to be transferred in small extracellular vesicles (sEV).

Overexpression of *Spp1* in neonatal chromosome 14 congenic and SHRSP strains preceded development of increased cardiac mass and onset of hypertension. The congenic strategy identified *Spp1* as a positional and functional candidate gene determining increased LVMI in the SHRSP model of human cardiovascular disease.

## 1 Introduction

Cardiovascular disease (CVD) remains a leading cause of morbidity and mortality worldwide (1). Left ventricular mass index (LVMI) is a strong, independent predictor of cardiovascular events and death (2), however mechanisms leading to a physiological enlargement of the LV are distinct from those that are associated with pathological LVH (3). Although LV remodelling is initially an adaptive response increased load, pathological LV hypertrophy (LVH) is associated with adverse cardiovascular outcomes in population-based studies (4–6).

The development of pathological LVH is influenced by multiple factors, including the severity and duration of pressure overload, age, sex, ethnicity, and adverse metabolic characteristics, such as increased circulating glucose, cholesterol, and triglyceride levels (6–8). Nevertheless, these clinical and haemodynamic factors explain only 50-70% of the observed inter-individual variation in LVMI (6). Evidence from twin and familial studies indicate that cardiac mass is highly heritable and influenced by numerous genetic loci, consistent with a polygenic architecture (9,10). These findings highlight the important contribution of genetic factors to pathological LV remodelling and underscore the need for genomic studies to unravel the genetic basis of this complex phenotype (11).

The identification of causal genes and pathways can be aided by experimental models of CVD that are genetically controlled and more easily modifiable. The Stroke Prone Spontaneously Hypertensive Rat (SHRSP) is a well characterised rat model of human hypertension and associated end organ damage. Compared to its closest genetic control, the Wistar Kyoto (WKY), SHRSP animals develop cardiac hypertrophy, increased stochastic incidence of stroke, and progressive renal disease alongside increased blood pressure in adulthood (12,13). Heart weight of inbred rat strains is also highly heritable, and partially independent from blood pressure (14).

Linkage of continuous traits by QTL analysis has identified more than 50 regions of the rat genome associated with structural and functional parameters of the heart. An F_2_ cross of the WKY and SHRSP previously identified a region of chromosome 14 independently associated with LVMI (15). The same study identified sex-specific linkages to blood pressure on rat chromosome 2, which were subsequently validated by the establishment of reciprocal chromosome 2 congenic strains on both WKY and SHRSP backgrounds (16). Transcriptome profiling, fine mapping of variants, and transgenic rescue identified glutathione S-transferase µ1 (*Gstm1)* as a novel positional and functional candidate gene in the control of blood pressure (17). The most recently published and largest genome wide association study (GWAS) for blood pressure control in 1 million individuals identified a single nucleotide polymorphism (SNP) upstream of the GSTM1 gene (rs36209093) associated with diastolic blood pressure (11).

Congenic strains enable refinement and validation of QTLs and facilitate the integration of genetic and transcriptomic data to identify causal genes in polygenetic traits (16–18). Correlation of positional variance with gene expression data can further implicate candidate genes causally associated with cardiac mass. We applied the speed congenic protocol to generate a chromosome 14 congenic strain from WKY (background) and SHRSP (donor) strains to confirm the genetic linkage between rat chromosome 14 and LVMI (15). We hypothesised that introgression of the SHRSP chromosome 14 region on the WKY background would result in an increase in congenic LVMI, independent of blood pressure. Further, structural alterations in the LV would be coupled to, and preceded by, differences in the cardiac transcriptome that could be utilised to finely map positional and functional candidate genes influencing LVMI.

The congenic strategy identified secreted phosphoprotein 1 (*Spp1),* also known as osteopontin, as consistently upregulated in SHRSP and congenic strain LV across all age groups. *In silico* analysis of the *Spp1* promoter region predicted variants in the SHRSP genome increase T-box factor binding, relevant for cardiac development and response to cardiac injury. *Spp1* overexpression was functionally assessed in the H9c2 cardiac cell model and was shown to significantly increase cell size. The transfer of *Spp1* in small extracellular vesicles (sEV) was also investigated in the hypertrophic effect of increased *Spp1* expression. We provide evidence that the LVMI-linked region of SHRSP chromosome 14 affects heart mass, diastolic function, and cardiac fibrosis. *Spp1* is therefore proposed to be a positional and functional candidate gene within the chromosome 14 congenic interval.

## 2 Methods

### Generation of Chromosome 14 Congenic Strains

Animal procedures were approved by the Home Office according to the Animals Scientific Procedures Act (1986) and subject to local ethics approval. Animals were kept at 21°C ambient temperature, on a 12-hour light/dark cycle with *ad libitum* access to chow diet and water. Inbred colonies of WKY and SHRSP have been maintained at the University of Glasgow since 1991 (WKY_Gla_, and SHRSP_Gla_). A ‘marker-assisted’ speed congenic approach was implemented to produce a congenic strain on WKY background, containing aregion of the SHRSP genome on chromosome 14, previously linked to LVMI by QTL analysis. The nomenclature of the strain, WKY.SPGla14a, depicts first the recipient (background, WKY) strain, then the donor strain (SP), and finally the colony origin (Gla), and the rat chromosome targeted for congenic generation (14a). The WKY.SPGla14a contains a 29Mbp region of SHRSP chromosome 14 genome between simple sequence length polymorphism (SSLP) markers D14Rat54 - D14Got41, which effectively maps to chr14:4,041,872-4,042,062 (+) and chr14:32,102,616-32,102,744 (+) on the mRatBN7.2 reference genome assembly.

### Blood Pressure and Cardiac Phenotyping of Parental and Congenic Strains

WKY, SHRSP, and WKY.SPGla14a strains were assessed at 3 timepoints representing early post-natal development (day 1-3 neonate), juvenile (5-weeks), and adult (12-20weeks). Male rats were used for 5- and 16-week assessments. Systolic and diastolic blood pressure was measured over an 8-week period using the Dataquest ART Telemetry System (Data Sciences International). Rats were implanted at 11-weeks of age and after 1-week recovery, probes were switched on and data were collected twice per week, for 8 weeks.

Transthoracic echocardiography was performed at 5 and 16 weeks using the Acuson Sequoia c512 ultrasound system with a 15-MHz linear array transducer to acquire M-mode images. Data across images were averaged and used to derive an estimate of LVM using a cubed model, which was corrected for body mass to generate individual LVMI. A terminal measure of LV diastolic and systolic function was made at 16-weeks using the Scisense pressure-volume (PV) system (Supplementary methods).

Neonate rats were euthanised by decapitation followed by exsanguination between post-natal days 1-3. Rats of 5- and 16-weeks of age were anaesthetised and euthanised with isofluorane (4%) followed by exsanguination. The heart was removed, cleaned and LV dissected. Soluble collagen was detected using the Sircol collagen assay kit. LV apex at 16-week was formalin-fixed and paraffin embedded for picrosirius red staining and quantification of *Spp1*/osteopontin (supplementary methods).

### Gene Expression Profiling and *in-silico* Transcription Factor Binding

Gene expression was assessed in the whole heart of 1-day neonate rats, and the LV of 5 and 16 week rats (N=3/strain) using the RatRef-12 v1.0 microarray chip (Illumina). Whole hearts from 1-day neonate animals were collected, weighted and snap frozen for RNA extraction. LV was snap frozen following dissection and weighting from 5 and 16 week rats. RNA was extracted from frozen heart/LV tissue using RNeasy kits according to the manufacturer’s protocol (Qiagen). Biotinylated amplified target cDNA was prepared and hybridised to the chip. Chip data was normalised, and differential expression analysis was performed in R using *limma-voom*(19). Following *voom* transformation and calculation of variance weight, *limma* was used to fit a linear model of weighted least squares for each gene. Contrasts with empirical Bayes smoothing were made between the SHRSP and WKY.SPGla14a strains *vs* WKY, at each time point (1-day neonate, 5 weeks, 16 weeks). Age comparisons (5 weeks *vs* neonate) were also made within in each strain. Raw expression data is accessible via GSE344388.

Differences in transcription factor binding were tested computationally using the MotifBreakR package (20) and SearchSeq function of TFBSTools (21). The 2024 version of the JASPAR Core database was used to generate the required position weight matrices (PWM) for transcription factors. In both analyses, 5000bp upstream of transcriptional start site (TSS) was used to determine potential *cis*-regulatory effects.

### Quantitative Reserve Transcription PCR

Taqman qRT-PCR was performed on the QuantStudio 12K Flex Real-Time PCR System (Thermo-Fisher, Scientific, UK) using cDNA prepared from RNA using SuperScript II and non-specific primers. cDNA (2µL) was combined in a multiplex reaction with a master-mix containing FAM-conjugated probes for (*Spp1* (Rn00681031_m1), and a VIC-conjugated probe for *B2m* (Rn00560865_m1). Biological replicates were performed in triplicate and average cycle threshold values (Ct) were analysed using the 2 method (22). The expression of target genes was normalised to *B2m*, and quantified relative to expression in the WKY.

### Cell Culture, Extracellular Vesicle Isolation and Assessment of Cell Size

cDNA templates for the *Spp1* gene from WKY and SHRSP genomes were incorporated into cloning vectors and isolated plasmid DNA was used in overexpression assays. H9c2(2–1) cells (ATCC CTRL-1446) were transfected for 48 hours with angiotensin-II (AngII), or plasmids containing pcDNA (plasmid control) or *Spp1* using lipofectamine. Cell lysates and conditioned media were submitted for liquid chromatography mass spectrometry (LC-MS) for untargeted metabolomic analysis (supplementary methods).

Cell sizing assays were carried out 48 hours following transient transfection with either; Ang-II, plasmids containing *Spp1*, co-cultured with conditioned medium from *Spp1* transfected cells (CCM), or co-culture with sEV isolated from transfected cells (supplementary methods). H9c2 cells were fixed in 4% PFA and stained using 0.5% crystal violet stain. Plates were visualised using an EVOS Core Cell Imaging System (ThermoFisher Scientific, UK). Cell length was measured using the straight-line tool in ImageJ, along the long axis of each cell. ‘Cell size’ describes this measure of H9c2 cell elongation herein.

EV uptake was blocked in H9c2 cells by adding 60µM dynasore, delivered in 1% DMSO to co-culture experiments. Co-culture in the presence of 1% DMSO was used as a vehicle controls. After 48 hours of culture with sEV ± dynasore, cells were fixed, stained and imaged as described above. RNA was extracted from *n*=3 EV suspensions using miRNeasy kit (Qiagen) as per the manufacturer’s instructions. RNA quality was assessed using the Agilent BioAnalyzer at Glasgow Polyomics who performed library preparation, amplification, and RNA sequencing. Reads in raw .fastq files were trimmed and filtered by Glasgow Polyomics using standard FastQC and fastp. Reads and were aligned and quantified using Kallisto (23) and abundance files read into R using Tximport (supplementary methods).

### Statistical Analysis

Echocardiographic and H9c2 cell image analysis was conducted in ImageJ. Downstream processing and statistical analysis of all data was performed in R (R v4.3 or v4.4). Pairwise comparisons were made following one-way ANOVA. Dunnett’s test was used to compare *in-vivo* data to the WKY as control and other *post-hoc* comparisons were made using Tukey. When the equality of variance assumption did not hold, Welch’s ANOVA and equivalent pairwise comparisons were used (Games-Howell). When only two groups were compared, a two-sample t-test was used. A *p*-value of <0.05 was considered statistically significant.

## 3 Results

### Rat Chromosome 14 Independently Influences Cardiac Structure and Function

The WKY.SPGla14a congenic strain was produced by marker-assisted selection of offspring between markers D14Rat54 and D14Got41 (Figure 1a). The congenic strain was fixed by brother-sister mating and confirmed by genotyping.

**Figure 1.**
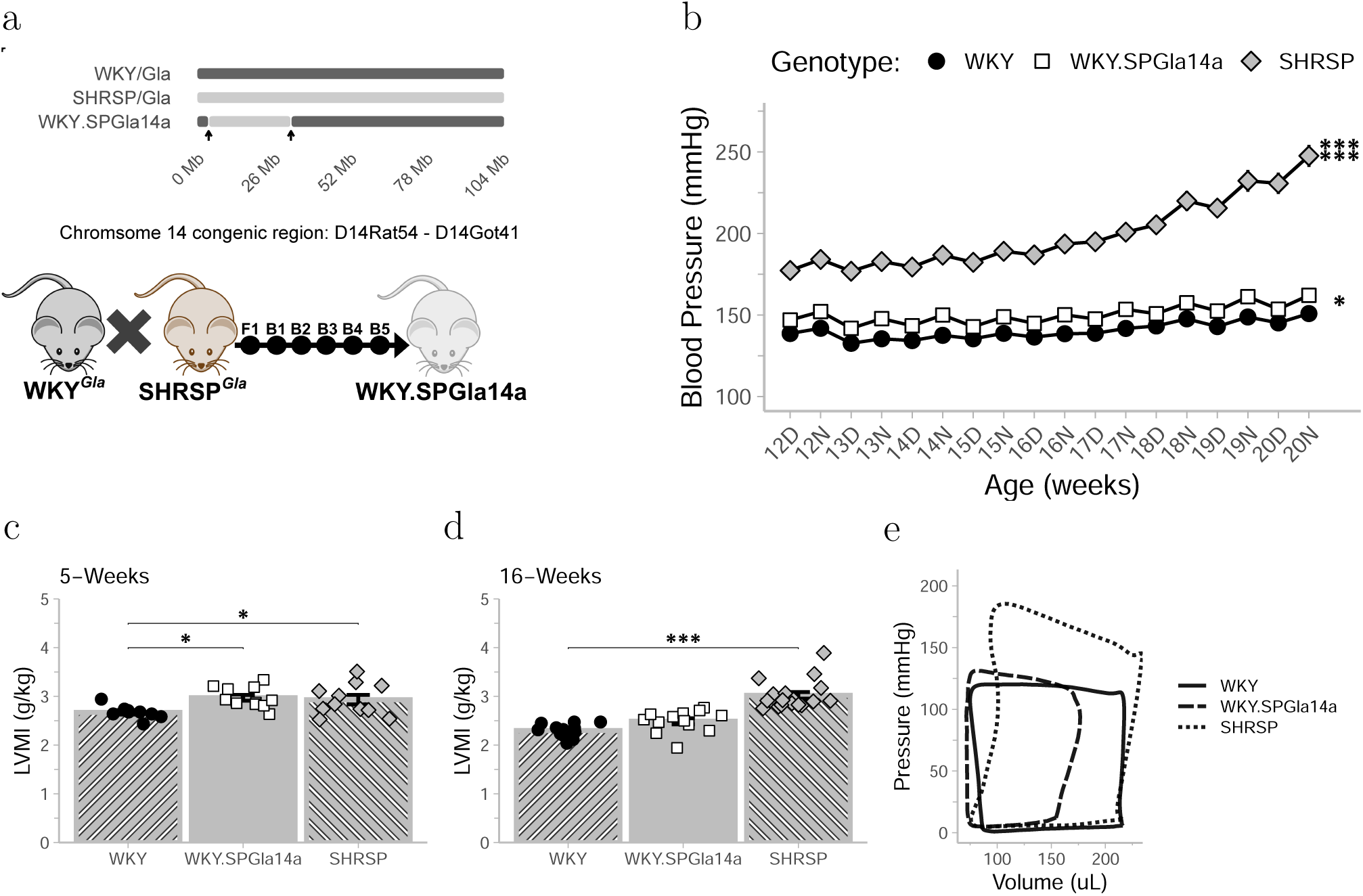
(a) Graphical representation of congenic region in WKY.SPGla4a and speed congenic protocol adopted for strain generation. (b) Daytime (D) and night-time (N) average blood pressure recorded by radiotelemetry over a 9-week period in the SHRSP (*n*= 13), WKY.SPGla4a (*n*= 8), and WKY (*n*= 7) strains. Welch-ANOVA of area under curve (AUC, F(2,16.29) = 82.6, *p*= *<*0.0001) was followed by Games-Howell pairwise comparisons. (c-d) Left ventricular mass index (LVMI, g/kg body mass) estimated by echocardiography in 5- and 16-week male WKY (*n*_5_= 9, *n*_16_= 12), SHRSP (*n*_5_= 11, *n*_16_= 18) WKY.SPGla14 (*n*_5_= 12, *n*_16_= 13) strains. Significant pairwise comparisons following one-way ANOVA (F_5-week_(2,29)= 4.62, *p*= 0.018, F_16-week_(2,40)= 38.36, *p*<0.0001) are displayed in plot area. Dunnetts test was used to perform pairwise comparisons, using WKY as the reference strain. (d) Example Pressure-Volume recording from WKY (*n*= 7), WKY.SPGla14a (*n*= 5) and SHRSP (*n*= 7) strains. All data is shown as mean *±* SEM. \**p<*0.05, \*\**p<*0.01, \*\*\**p<*0.001

The adult SHRSP showed significantly higher blood pressure than both the WKY (AUC_diff_=497, *p*<0.001) and WKY.SPGla14a (AUC_diff_=412, *p*<0.001), at all time-points measured between 12 and 20-weeks of age (Figure 1b). Blood pressure in the WKY.SPGla14a strain was marginally but significantly increased compared to the WKY (AUC_diff_=84.6, *p*=0.045).

At 5 weeks, both SHRSP (LVMI_diff_=0.26g/kg, *p*=0.038) and WKY.SPGla14a (LVMI_diff_=0.30g/kg, *p*=0.014) had significantly increased LVMI compared to WKY (Figure 1c), which persisted to 16-weeks in SHRSP (Figure 1d, LVMI_diff_=0.72g/kg, *p*<0.001).

Diastolic function, as assessed by the stiffness constant end-diastolic pressure-volume relationship (β), was significantly increased in SHRSP compared to WKY (Table 1). β was increased in WKY.SPGla14a compared to WKY, although did not reach significance in *post-hoc* testing. Representative traces indicate the increase in stiffness with an increase in the slope between the end of systole and beginning of diastole (Figure 1e). In both SHRSP and WKY.SPGla14a, end systolic pressure (ESP) was significantly increased compared to WKY. Both the maximal rate of rise in pressure (dP/dt_max_), and the maximal rate of fall in pressure (dP/dt_min_) were significantly increased in SHRSP compared to WKY. These differences are reflected in representative P-V loops from each strain (Figure 1e). SV and cardiac output (CO) were reduced non-significantly in the SHRSP compared to the WKY. The reduced SV in WKY.SPGla14a contributed to a significant decrease in CO compared to WKY (Table 1, Figure 1e).

**Table 1.** Haemodynamic PV data from the background and chromosome 14 congenic strains.

|  | WKY | WKY.SPGLa14a | SHRSP | *p*-value vs WKY |  |
| --- | --- | --- | --- | --- | --- |
|  | *n* = 7 | *n* = 5 | *n* = 7 | WKY.SPGLa14a | SHRSP |
| **HR** | 325.771<br>(±23.737) | 336.572 (±3.982) | 350.884<br>(±31.302) | 0.70 | 0.12 |
| **ESP** | 125.157<br>(±8.104) | 139.905 (±3.451) | 185.562<br>(±10.488) | **0.015** | **<0.001** |
| **EDP** | 7.650 (±2.432) | 11.145 (±2.436) | 11.209 (±4.158) | 0.14 | 0.10 |
| **dP.dt<sub>max</sub>** | 7,796.4<br>(±1,041.1) | 8,042.9 (±498.1) | 11,092.7<br>(±1,325.9) | 0.90 | **<0.001** |
| **dP.dt<sub>min</sub>** | 8,085 (±720.4) | 8,243 (±498.1) | 11,309.4<br>(±1,267.8) | >0.90 | **<0.001** |
| **Tau (τ)** | 10.372 (±0.960) | 10.356 (±0.896) | 9.706 (±0.688) | >0.90 | 0.30 |
| **EDPVR** | 0.011 (±0.010) | 0.034 (±0.014) | 0.034 (±0.021) | 0.058 | **0.048** |
| **ESV** | 116.058<br>(±29.219) | 104.613 (±5.866) | 172.561<br>(±40.174) | 0.80 | **0.013** |
| **EDV** | 259.006<br>(±49.044) | 202.902 (±22.293) | 296.075<br>(±69.252) | 0.20 | 0.40 |
| **SV** | 142.941<br>(±37.629) | 98.292 (±17.595) | 123.510<br>(±49.847) | 0.20 | 0.60 |
| **CO (mL/min)** | 46.506 (±9.162) | 33.073 (±6.285) | 38.075<br>(±10.248) | **0.046** | 0.20 |
| **EF** | 51.981<br>(±10.236) | 48.205 (±3.399) | 40.986 (±8.916) | 0.70 | 0.058 |

HR = heart rate; ESP = end-systolic pressure; EDP = end-diastolic pressure; dP/dt<sub>max</sub> = maximal rate of rise of pressure; -dP/dt<sub>min</sub> = maximal rate of fall in pressure; Tau (τ) = relaxation time constant; EDPVR = end-diastolic pressure-volume relationship stiffness constant. ESV = end-systolic volume; EDV = end-diastolic volume; SV = stroke volume; EF = ejection fraction; CO = cardiac output. Values are expressed as mean (SD). Dunnett's pairwise comparison to WKY was made following one-way ANOVA.

Interstitial fibrosis was significantly influenced by genotype (F(2,5.29) = 8.3, *p*=0.023, Figure 2a). Positive area was increased in the LV of WKY.SPGla14a (%diff=2.84, *p*=0.034) compared to WKY (Figure 2a & Figure 2d). The comparison between SHRSP and WKY did not reach significance after adjustment for multiple testing. Histological assessment of perivascular fibrosis did not reach significance by 16-weeks of age (Figure 2b), however a trend toward increased % positive staining was present in WKY.SPGla14a and SHRSP compared to WKY. LV collagen content (µg/µL) was significantly increased in WKY.SPGla14a and SHRSP strains compared to WKY at 16 weeks of age (Figure 2c).

**Figure 2.**
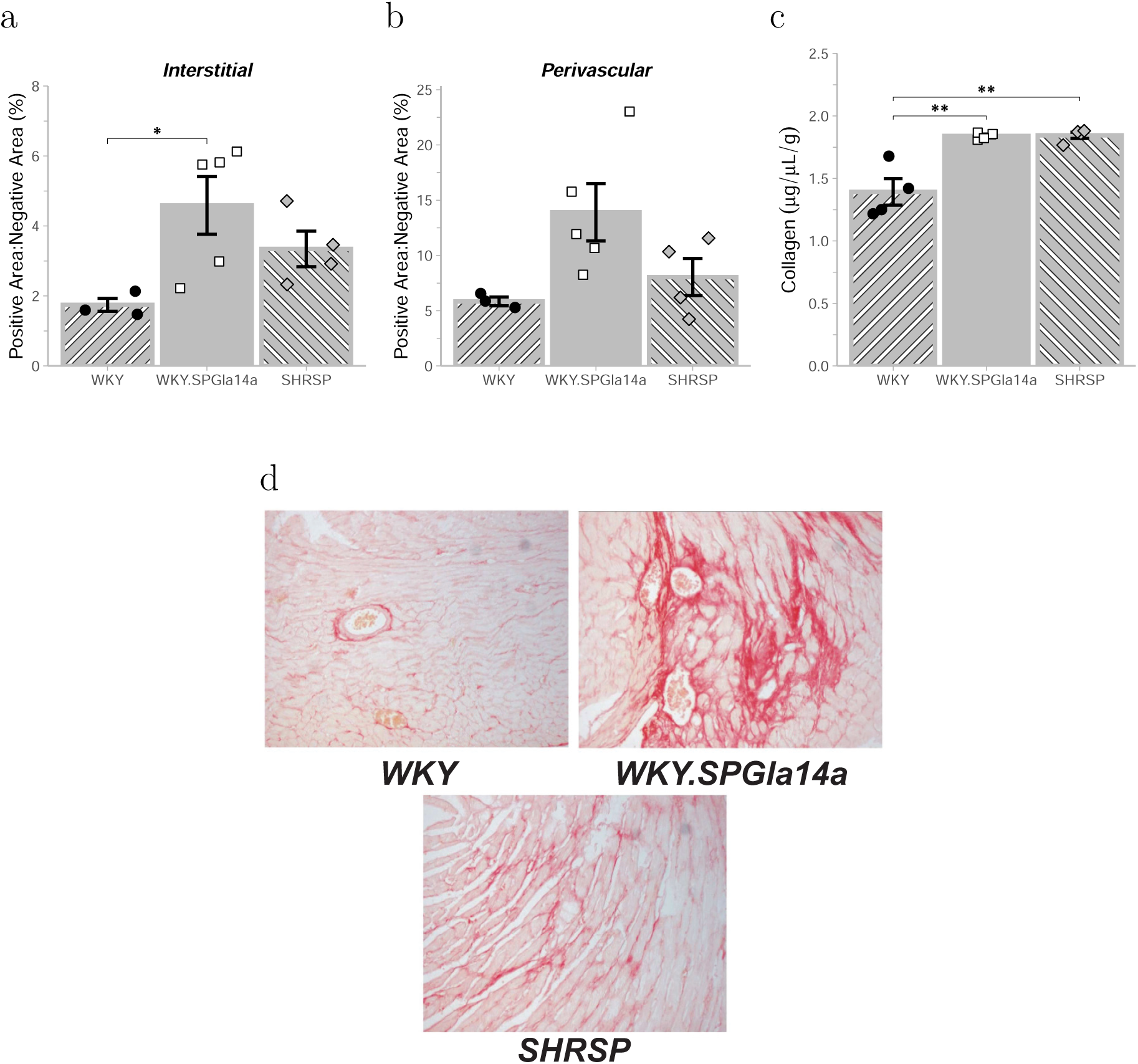
Assessment of LV fibrosis in WKY, WKY.SPGla14a and SHRSP strains (*n*= 3-4/group). (a) Interstitial and (b) perivascular fibrosis was measured in picrosirius red stained LV sections from WKY (*n*= 3), WKY.SPGla14a (*n*= 5), and SHRSP (*n*= 4, F_interstitial_ (2,5.29)= 8.3, *p*= 0.023, F_perivascular_ (2,4.77)= 4.77, *p*= 0.07). (c) Collagen content was measured and quantified in LV tissue using colorimetric assay (*n*= 4/group). Significant one-way ANOVA (F(2,9) = 5.7, *p*= 0.025) was followed by Dunnett’s test and significant pairwise comparisons are displayed in plot area. (d) Representative images of picrosirius red staining in the LV. All data is shown as mean *±* SEM. \**p<*0.05, \*\**p<*0.01, \*\*\**p<*0.001

### Changes in Cardiac Transcriptome Precedes Development of Increased LVMI in SHRSP and WKY.SPGla14a Strains

Cardiac weight (heart weight mass index, HWMI) was decreased in 1-day neonates of SHRSP (HWMI_diff_=-0.55, *p*=0.031) and WKY.SPGla14a (HWMI_diff_=-0.56, *p*=0.038) strains compared to age-matched WKY (Figure 3a).

**Figure 3.**
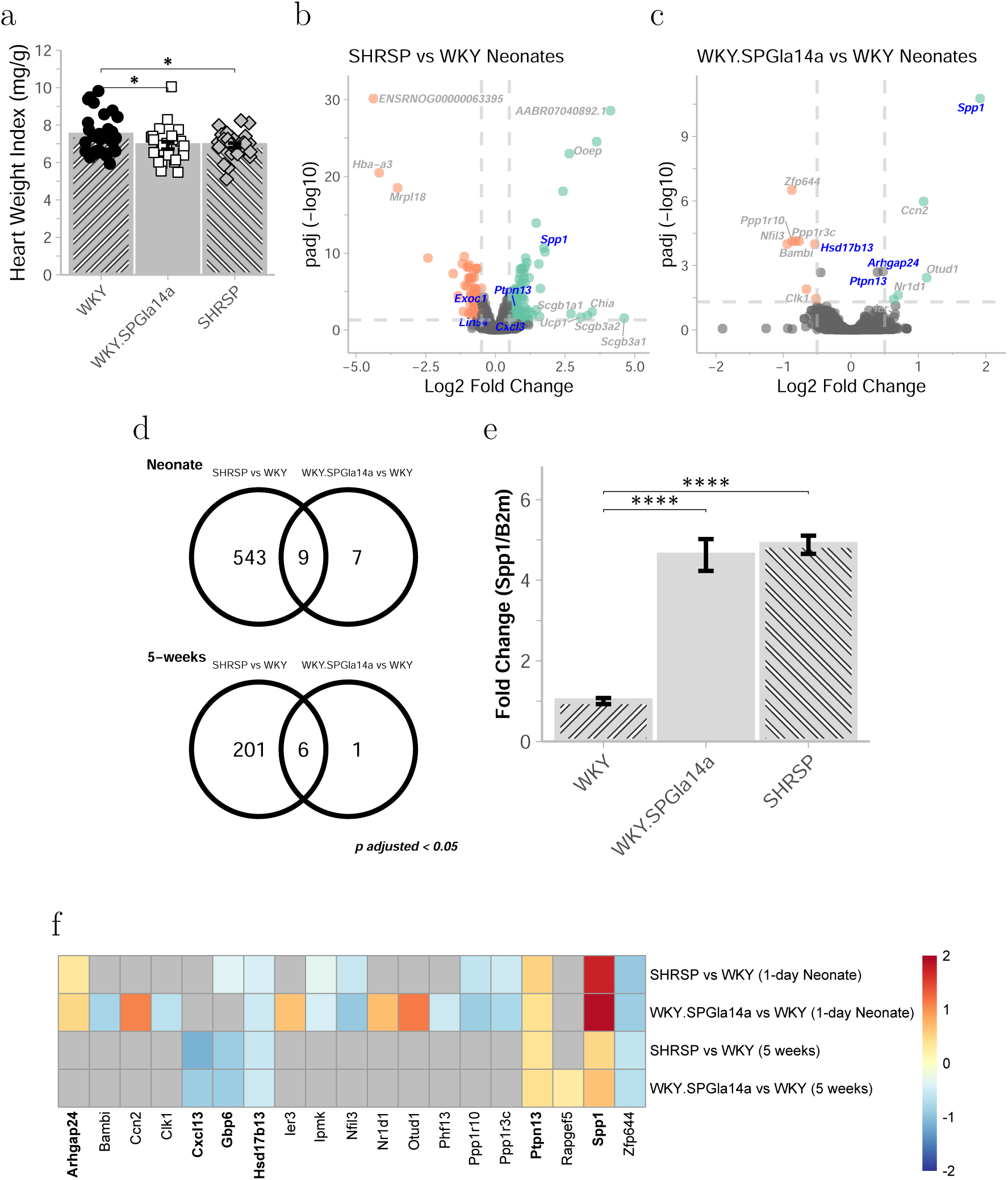
(a) heart to body weight index of 1-3 day neonate hearts, Dunnett’s pairwise comparisons of SHRSP (*n*= 30) and WKY.SPGla14a (*n*= 25) to WKY (*n*= 33) are displayed in plot area (F(2, 85)= 3.996, *p*= 0.022). (b-c) volcano plot of differential gene expression analysis in 1-day neonates (*n*= 4/group), genes with FDR*<*0.05 and log2 fold change >*±*0.5 are coloured orange (negative fold change) and green (positive fold change). Genes with the highest log fold change are labelled, blue text signifies genes located within the chromosome 14 congenic region introgressed into the WKY.SPGla14a. (d) Venn diagram of genes with significant differential expression (*p_adj_*<0.05) between SHRSP or WKY.SPGla14a and WKY at neonate (top) and 5 week (bottom) timepoints. (e) Taqman qRT-PCR validation of *Spp1* in neonate hearts (*n*= 4/group). Tukey posthoc comparisons following significant one-way ANOVA (F*_Spp1_* (2,9)= 161.21, *p*<0.0001) are shown on the plot area. (f) Heatmap of log fold change in 19 genes showing significantly different expression WKY.SPGla14a at neonate and/or 5 weeks (one or both time-points). Genes with a non-significant difference in expression are coloured grey. \**p≤*0.05, \*\**p≤*0.01, \*\*\**p≤*0.001, \*\*\*\**p≤*0.0001.

As expected, comparison of cardiac gene expression between SHRSP and WKY.SPGla14a, relative to WKY showed the number of genes with significantly different expression was substantially lower in the WKY.SPGla14a *vs* WKY, than the SHRSP *vs* WKY comparison, at all timepoints (Figure 3b-d, & Figure S1). However, a considerable proportion of WKY.SPGla14a *vs* WKY changes were shared with SHRSP *vs* WKY (Figure 3d, Figure S1f). The cardiac transcriptome of SHRSP and WKY.SPGla14a shared differences in the expression of 9 genes in 1-day neonates, and 6 genes in the 5-week comparisons, *vs* WKY. These shared differences were in 11 unique genes and showed the same direction of change was the same across both comparisons (Figure 3f). In 1-day neonate hearts, *Arhgap24*, *Ptpn13*, and *Spp1* were upregulated in the SHRSP and WKY.SPGla 14a (Figure 3f). These genes are located within the chromosome 14 region introgressed into the WKY.SPGla14a from the SHRSP. Also, within the congenic region, *Hsd17b13* and *Zfp644* were significantly down-regulated in SHRSP and WKY.SPGla14a compared to WKY (Figure 3f). The upregulation of *Spp1* in SHRSP and WKY.SPGla14a neonate heart was validated by qRT-PCR (Figure 3e). *Spp1* encodes the osteopontin (OPN) protein. There was a small increase in the expression of OPN in 16-week WKY.SPGla14a and SHRSP hearts (Figure S2). Due to the established association between *Spp1* and cardiovascular diseases, investigation focused on the role of *Spp1* in LVH development.

### Transcription factor binding analysis *in-silico* predict differences in binding upstream of *Spp1* transcription start site

The -5kb region upstream of *Spp1* contains a number of small variants in the WKY compared to the SHRSP (Figure S3a). SearchSeq analysis of 5kb upstream of the *Spp1* transcription start site (TSS) predicted the sequence contained >800 potential TF binding sites, most of which were found in both strains (Figure 4a). There were 3 TFs predicted to bind specifically in the SHRSP promoter sequence (Figure 4c), including T-box factors within 100bp of the TSS. In the overlapping hits, delta scores between the WKY and SHRSP were calculated for each TF motif ID. SearchSeq predicted higher binding affinity of the zinc-finger TF Wilms tumour 1 (Wt1) in the WKY compared to the SHRSP (Figure 4b). MotifBreakR analysis was used as a second *in-silico* investigation of TF binding, where variant data from alignments of the WKY and SHRSP to the mRatBN7.2 is used to predict variants impact on TF binding to motifs. Motifs that were affected by common variants, shared between the Wistar derived WKY/SHRSP and the mRatBN7 were removed. MotifBreakR predicted that variants in the WKY genome could significantly increase Wt1 binding (Figure 4d), which was in agreement with the SearchSeq prediction. Within -1kb of *Spp1* TSS, the WKY has a T/C allele disrupting the CCTCCCCCAC motif (Figure 4e). From the family of T-box factors uniquely identified in SearchSeq analysis, the TBX15 and TBX18 TFs were predicted to be affected by a C/T SNP, increasing affinity to the common AGGTGGGA motif (Figure 4d & Figure 4f).

**Figure 4.**
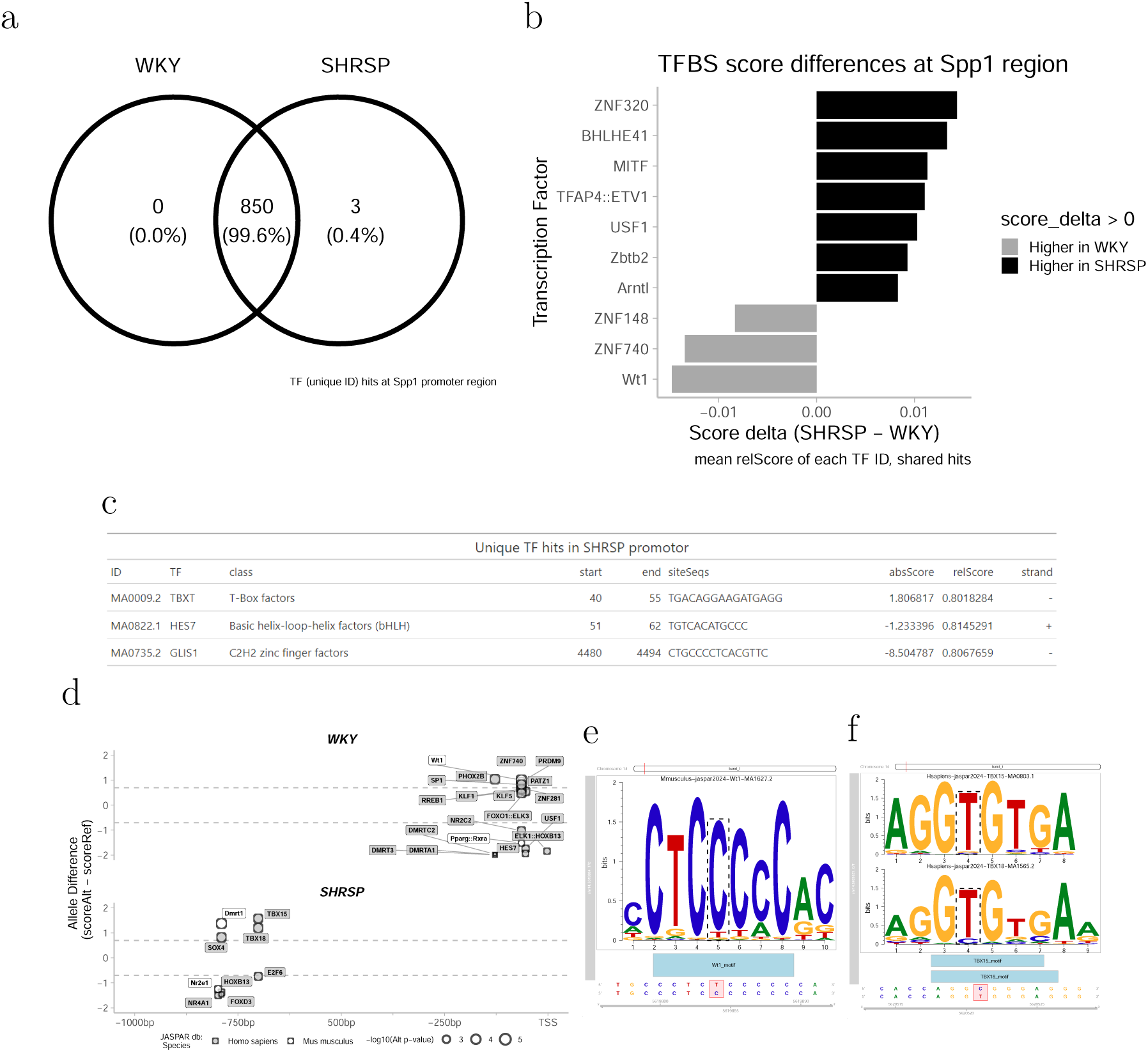
Computational transcription factor analysis of 5kb upstream of *Spp1* transcriptional start site (TSS) in the WKY and SHRSP specific genome sequence. (a) Number of TF ID hits with a relative score >0.8 from SearchSeq analysis in the *Spp1* +5kb promoter region. (b) TF were grouped by ID and an average score was computed in each genome. Delta score SHRSP-WKY is the comparison of the mean relative TF binding score between the genomes. Black bars show increased relative binding score in the SHRSP and grey bars show decreased relative score in the SHRSP compared to the WKY. (c) Unique SearchSeq hits in SHRSP *Spp1* promoter region. (d) Short variants that are predicted to increase or decrease TF binding as predicted by MotifBreakR are shown as distance from TSS. Unique SNPs and INDELs in WKY and SHRSP -2kb from TSS are displayed, labelled with the TF name. Size of point depicts adjusted *p*-value as produced by MotifBreakR. (e) Wt1 motif and SNP promoting binding in WKY genome (f) TBX15/TBX18 motif and SNP promoting binding in SHRSP genome.

### *Spp1* activation is independent of AngII stimulation and increases H9c2 cell size

The amino acid sequence of WKY and SHRSP *Spp1* cDNA was aligned by multiple sequence alignment (Figure 5a). A SNP in the SHRSP cDNA results in an isoleucine-leucine amino acid switch in the SHRSP *Spp1* protein. H9c2 cells were directly transfected with plasmid DNA containing *Spp1* derived from both WKY and SHRSP strains to induce over-expression. *Spp1* derived from both strains significantly increased H9c2 cell size after 48 hours (Figure 5b). The magnitude of effect was unaffected by the amino acid change between the strains and similar to that induced by known hypertrophic agent, AngII (Figure 5b). Assessment of *Spp1* expression in H9c2 cells following Ang-II or *Spp1* stimulation indicated Ang-II did not increase *Spp1* expression in H9c2 cells relative to control cells (Figure 5c). In contrast, transient overexpression of *Spp1* caused an increase in *Spp1* mRNA expression 48 hours following transfection (Figure 5c).

**Figure 5.**
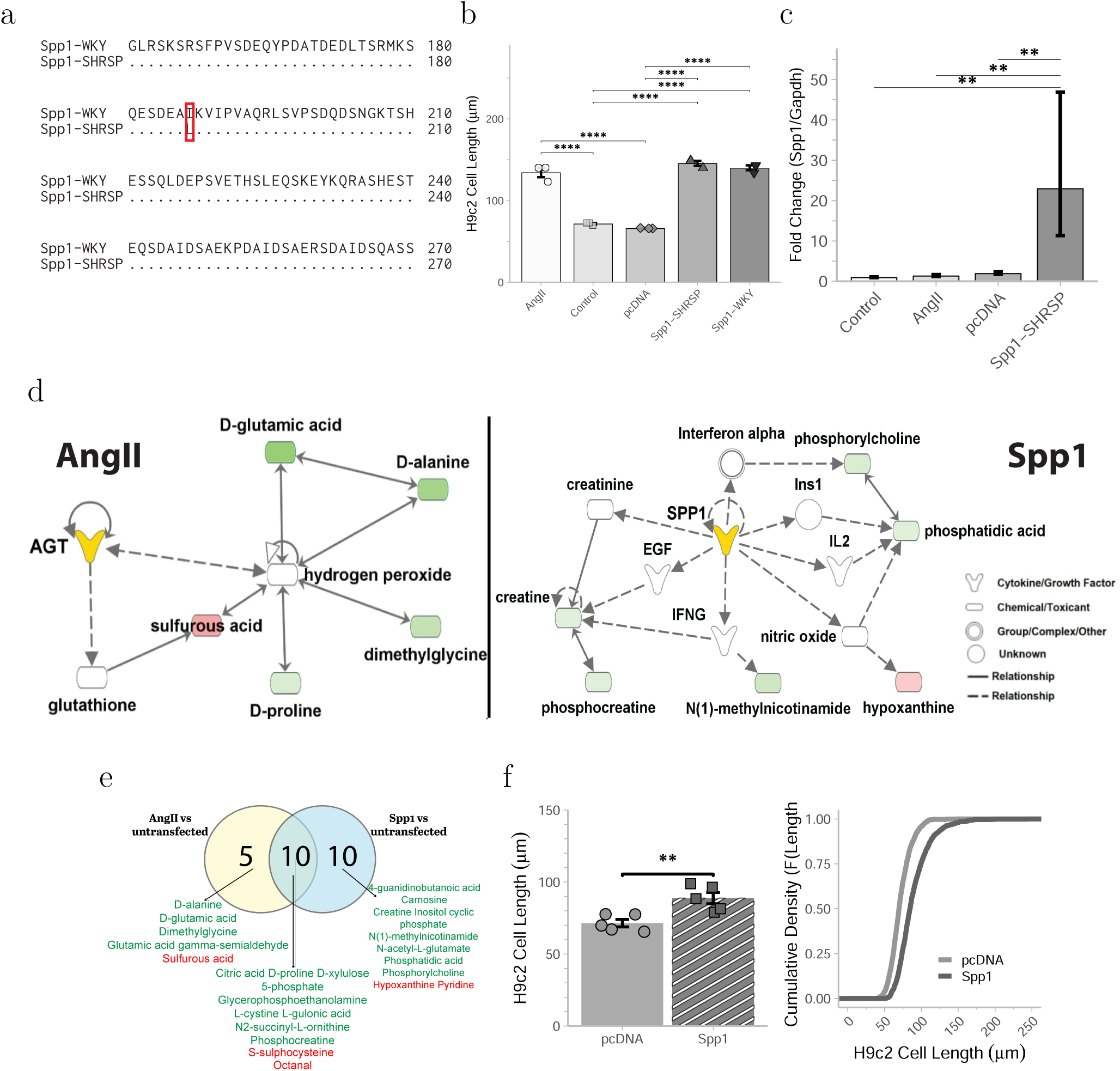
(a) Residues 151-270 of pairwise sequence alignment of *Spp1* protein derived from the SHRSP and WKY *Spp1* cDNA sequence. Matching residues denoted by *·* symbol. Non-matching residue at position 186 is labelled with amino-acid symbol. (b) Mean cell length of H9c2 cells 48 hours after; treatment with AngII, or transfection of plasmid containing *Spp1* cDNA derived from the WKY, SHRSP, or pcDNA. No transfection/treatment was included as a baseline control (*n*= 3/condition, F(4,10)= 148.9, *p<*0.0001).(c) H9c2 expression of *Spp1* following stimulation with AngII, no transfection control, pcDNA plasmid or plasmid containing SHRSP-*Spp1* (*n*= 3 /condition, F(3,8)= 14.65, *p*= 0.001).(d) Representative figure generated in Ingenuity Pathway Analysis (IPA) of relationships between metabolites identified in the conditioned media from AngII or *Spp1* treated cells. (e) Common and treatment specific changes (FDR*≤*0.05) to the metabolome of H9c2 cells following *Spp1* overexpression or AngII stimulation are displayed in venn diagram and annotated. Metabolites in green were downregulated, whilst red indicates upregulation following treatment. (f) Mean cell length (left) and Empirical Cumulative Distribution Function (right) of naïve H9c2 48 hours following co-incubation with conditioned cell media collected from H9c2 cells transfected with pcDNA or *Spp1* (*n*= 5 per condition, t(7.21)= -4.46, *p*= 0.003)). All data is displayed as mean*±*SEM. \**p≤*0.05, \*\**p≤*0.01, \*\*\**p≤*0.001, \*\*\*\**p≤*0.0001.

In untargeted LS-MS metabolomics, cells clustered according to transfection condition (Figure S3bc) and identified 25 differentially expressed small molecules that could be reliably annotated in Ingenuity Pathway Analysis (IPA). Metabolites associated with *Spp1* expression were inflammatory cytokines and growth factors, whilst Ang-II transfection was associated with chemical toxicants linked to oxidative stress (Figure 5d). *Spp1* transfection also altered cell metabolism to a greater extent than Ang-II, exemplified by the greater number of metabolites differentially expressed in the *Spp1* condition (Figure 5e).

### Excess *Spp1* is released from H9c2 cells in extracellular vesicles

Adding pre-conditioned media from cells transfected with *Spp1* as the growth medium to naïve H9c2 cells resulted in an equivalent increase in H9c2 cell size after 48-hours (Figure 5f). sEV isolated from conditioned media of *Spp1* transfected H9c2 cells was characterised by nanoparticle tracking analysis. The size and concentration of sEV released by H9c2 cells was not altered by *Spp1* transfection (Figure 6a-b). Culturing H9c2 cells with sEV isolated from *Spp1* treated cells caused naïve H9c2 cell size to increase compared to untreated cells, and compared to co-incubation with sEV isolated from pcDNA transfected cells (Figure 6c).

**Figure 6.**
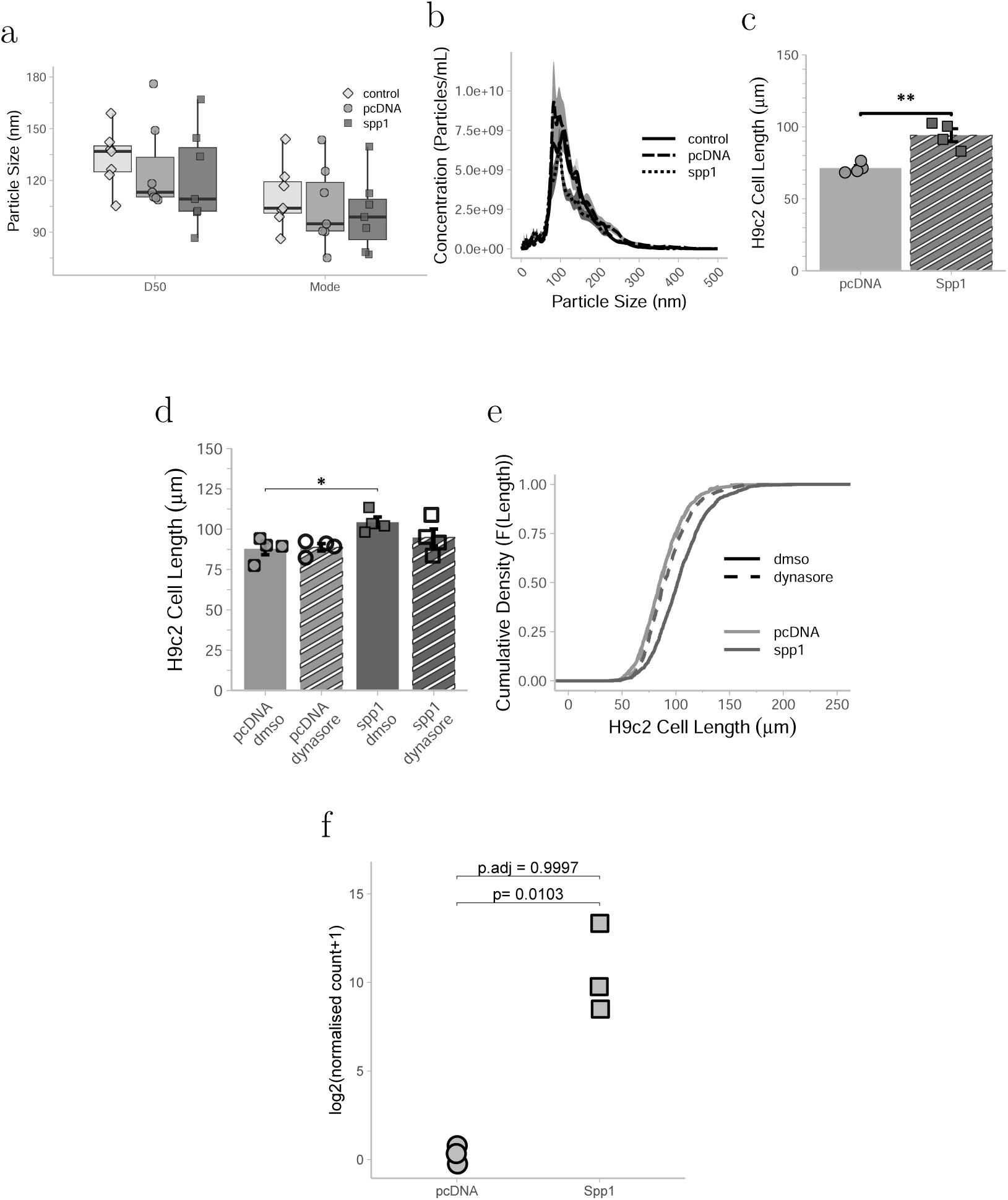
Nanoparticle tracking analysis of sEV isolated from H9c2 cells after 48 hours of pcDNA or *Spp1* transfection and no transfection controls. (a) Average particle size (D50 and Mode) is shown per transfection experiment. (b) Particle concentration is displayed in 0.5nm-increment bins (*n*= 7/condition). (c) Mean cell length of naïve H9c2 48 hours following co-incubation with small extracellular vesicles (sEV) collected from H9c2 cells transfected with pcDNA or *Spp1* (*n*= 4/condition, t(3.87)= -4.71, *p*= 0.01). (d) Mean cell length of navïe H9c2 cells (*n*= 4/condition, F(3,12)= 4.06, *p*= 0.033) and (e) Empirical Cumulative Distribution Function of cell populations 48 hours following co-incubation with vehicle control (DMSO, solid line) or Dynasore (dashed line) and isolated sEV from pcDNA or *Spp1* transfected cells. Comparisons was compared using 1-way ANOVA with Tukey HSD *post-hoc* comparisons displayed on the plot area. (f) Log2 normalised counts of *Spp1* from bulk RNA sequencing of sEV derived from pcDNA or *Spp1* transfected H9c2 cells (*p*= 0.01, *p_adj_*= 0.99). Data are displayed as mean*±*SEM; \**p≤*0.05, \*\**p≤*0.01, \*\*\**p≤*0.001, \*\*\*\**p≤*0.0001.

Cellular uptake of sEV can be blocked by co-incubation with the dynamin inhibitor dynasore. The addition of 60µM dynasore or 1% DMSO (vehicle control) did not change the effect of co-culture with pcDNA-associated sEV. In contrast, treatment of H9c2 cells with sEV isolated from *Spp1* treated cells resulted in a significant increase in H9c2 cell size in 1% DMSO (Figure 6d). This effect was reduced when 60µM dynasore was added, resulting in a left-ward shift of the ECDF curve (Figure 6e).

RNA sequencing of sEV isolated following transfection with pcDNA or *Spp1* could not reliably be used to build the DESeq2 model after filtering genes for low expression (Figure S4a-c) and high variability across the sEV samples did not produce any statistically significant differences in gene expression (*p*-adjusted=<0.05). However, sEV are not expected to contain the same diversity of mRNA found within the cells or tissues they are isolated from, which is currently unaccounted for in standard RNA sequencing analyses (Figure S4). As such, analysis was limited to determining whether *Spp1* could be reliably detected within sEV. Of the 200 genes with the greatest average normalised count across conditions, 161 were shared between sEV derived from pcDNA or *Spp1* transfected cells. *Spp1* mRNA was detectable in sEV isolated from H9c2s following *Spp1* transfection, but negligible in pcDNA-derived sEV (Figure S4f-S4g). Log2 normalised counts of *Spp1* within EVs from transfected cells were around 10 times higher than in sEV isolated from pcDNA transfected cells, but the difference was not significant after adjustment in DESeq2 (Figure 6f).

## 4 Discussion

Using a congenic strategy, the resulting WKY.SPGla14a strain has confirmed the positional genetic linkage previously identified between a region of rat chromosome 14 and LVMI (15). The SHRSP donor region is demonstrated to be a causal genetic determinant of cardiac mass, with WKY.SPGla14a rats displaying reduced postnatal cardiac mass and subsequent catch-up growth, resulting in increased LVMI by 5 weeks of age. This pathological “catch-up” growth of the heart is similarly reported in the hypertrophic heart rat model (24).

Importantly, introgression of the SHRSP chromosome 14 interval onto a normotensive background partially dissociated cardiac remodelling from increased blood pressure. Despite modest differences in blood pressure between WKY.SPGla14a rats and WKY controls, the WKY.SPGla14a developed structural and functional cardiac abnormalities similar to the SHRSP. By 16 weeks of age, both WKY.SPGla14a and SHRSP rats exhibited evidence of myocardial fibrosis, which was associated with increased ventricular stiffness, as assessed by pressure-volume loop analysis. Fibrosis and diastolic dysfunction were increased to a comparable extent in both strains relative to WKY controls, even with the more moderate blood pressure phenotype in WKY.SPGla14a. Together, these findings suggest that the chromosome 14 locus contributes directly to adverse cardiac remodelling, enabling partial dissociation of cardiac and hypertensive phenotypes.

The SHRSP chromosome 14 region was associated with upregulation of *Spp1*, located within the congenic interval of the WKY.SPGla14a. *Spp1* encodes osteopontin, which is a non-collagenous extracellular matrix protein in the highly conserved, small integrin binding ligand N-linked glycoprotein family (25). Increased *Spp1* expression is associated with heart failure, reduced diastolic function, and other cardiovascular phenotypes (26–28), whilst promoter variants of *Spp1* have been associated with diastolic function in a Japanese cohort of hypertensive adults (29). A transgenic mouse model of cardiomyocyte-specific *Spp1* overexpression develops a dilated cardiomyopathy-phenotype and dies prematurely from 8 weeks of age, despite normal cardiac development *in-utero* (30). In the same model, blocking cardiomyocyte overexpression of *Spp1* at 5 weeks, results in reduced mortality and restores survival to levels comparable with control mice (30). Thus, *Spp1* was proposed as a positional and functional candidate gene, supporting the genetic linkage between the chromosome 14 locus and cardiac mass.

Ang-II infusion, usually employed with the primary outcome of increasing blood pressure, also results in increased LVMI and cardiac fibrosis. Compared to wild type littermates, *Spp1* knock-out mice develop reduced fibrosis and lower cardiac mass following Ang-II infusion, despite comparable increases in blood pressure (31,32). In this study of genetically encoded hypertension and LV remodelling, we show the role of *Spp1* is potentially independent of the effect of Ang-II. Although Ang-II caused a significant increase in H9c2 cell size, this hypertrophic response was not accompanied by a change in *Spp1* expression. Consistent with the metabolomic profile of transfected H9c2 cells, Ang-II is reported to mediate cardiac hypertrophy through oxidative stress related pathways (33,34). In contrast, *Spp1* is most often described as an inflammatory mediator (35), functionally distinct from Ang-II.

A non-synonymous single nucleotide polymorphism (nsSNP), resulting in a single amino acid change, exists between WKY and SHRSP *Spp1* gene. Overexpression of both WKY and SHRSP forms of *Spp1* caused an increase in H9c2 cell size, supporting the hypothesis that continued overexpression of *Spp1* in SHRSP and WKY.SPGla14a strains causes observed increased LVMI, rather than altered function of the protein due to the nsSNP. *Spp1* expression is low in healthy cardiac muscle (32), however models of LVH in the rat (26,27,36) and mouse (32,37) indicate expression of *Spp1* is increased following pathological remodelling of the heart. In the myocardium, WKY.SPGla14a and SHRSP over-express *Spp1* during neonatal cardiac development, prior to the development of LVH, with elevated expression persisting until 5 weeks of age when LVMI is measurably increased.

*Cis*-acting regulatory pathways are likely contributing to this genetic effect, where the SHRSP genome reflects a risk profile. Variants within the SHRSP *Spp1* promoter are predicted to promote binding of t-box factors, TBX15 and TBX18, important for organogenesis, including development of the heart (38). Over-expression of *Tbx18* during mouse development alters the transcriptome, enriching for “hypertrophic/dilated cardiomyopathy” pathways (39), and causes development of cardiac fibrosis (40). Linage tracing coupled with single cell RNA sequencing showed *Tbx18*+ cells have increased *Spp1* expression in the brain (41). The WKY genome has been predicted to harbour a protective mutation increasing ability of Wt1 zinc-finger protein binding. Functionally opposing roles of the Tbx18 and Wt-1 transcription factors has been reported in the cardiac epithelium during mesenchymal transition of epicardial cells (42). Wt1-deletion in cardiomyocytes increases cardiac fibrosis in mice (43) and global Wt1 deletion increases *Spp1* expression (44), supporting a suppressive effect of Wt1 binding on *Spp1* expression (48). sEV or exosomes have been investigated as disease biomarkers and diagnostic tools in LV hypertrophy and progression to heart failure (45). These lipid-membrane bound vesicles contain cargoes of various bioactive molecules, including micro-RNAs, mRNA, protein, DNA, and lipids. EVs are produced by all cells that make up the heart and are involved in physiological and pathophysiological signalling (46). Rat cardiac fibroblasts mediate cardiomyocyte hypertrophy through release of exosomes containing the miRNA miR-21* (47). Isolated sEV derived from obese and hypertensive patients induce divergent responses in human induced pluripotent stem cell-derived cardiomyocytes (hiPSC-CMs) (48). H9c2 and hiPSC-CMs can alter EV size, shape, and cargo in response to ischaemic conditions (49). We demonstrate overexpression of *Spp1* does not change the morphology or number of EV released by H9c2 cells, however normalised counts of *Spp1* were much higher in sEV produced by cells stimulated to increase *Spp1* expression. The cellular model of *Spp1* overexpression in H9c2 cells support the hypothesis that release and uptake of *Spp1* mRNA within sEV is a potential mechanism by which *Spp1* mediates cellular hypertrophy.

Understanding genetic factors that determine LV mass is integral to improving therapeutic targets for LVH, which can be difficult to manage clinically. Sustained over-expression of *Spp1* in the heart from development into adulthood may promote increases in LVMI through molecular mechanisms that remain to be defined. It is unknown how the pathogenic SHRSP chromosome 14 region interacts with genetic and epigenetic factors to induce cardiac hypertrophy outside of increased *Spp1* expression, however *in-silico* analyses predict differences in zinc-finger and T-box transcription factor binding between the WKY and SHRSP genomes. Findings support a hypothesis that excess *Spp1* can be packaged into sEV, forming a positive feedback loop in which cells are activated to increase cell size. The congenic model was successfully applied to identify this positional and functional candidate gene involved in LVH pathology. Nevertheless, further studies are required to determine how *Spp1* interacts with the broader molecular networks dysregulated in WKY.SPGla14a and SHRSP rats, and whether these pathways converge with those implicated in human cardiovascular disease.

## 5 Perspectives

Translation of genome wide association studies (GWAS) and QTL analyses to molecular mechanisms can generate therapeutic targets. Here, SHRSP chromosome 14 is functionally linked with development of LV hypertrophy. Thus far, studies have assessed presence of *Spp1* after development of LVH. However, findings show *Spp1* represents an excellent positional and functional candidate gene for cardiac hypertrophy and associated fibrosis when over-expressed in cardiac muscle. Future investigations determining the mechanism by which *Spp1* influences cardiomyocytes and cardiac fibroblasts to promote hypertrophy and fibrosis is warranted, particularly exploring the role of small extracellular vesicles (sEV) and variants affecting TF binding in the *Spp1* promoter region.

## 6 Novelty and Relevance

What is new?

- A genetic link between a section of rat chromosome 14 and left ventricle growth has been confirmed, with *Spp1* emerging as a candidate gene of functional relevance.
- Overexpression of *Spp1* in cardiac cells can increase cell size and be transferred in small extracellular vesicles.

What is relevant?

- Left ventricular hypertrophy and hypertension share a causal relationship, however genetic determinants may be partially distinct, identifying additional targets beyond blood pressure control.

Clinical/Pathophysiological Implications?

- Chronic, genetically determined overexpression of *Spp1* is a potential target to reduce left ventricular hypertrophy in the context of hypertension.

## 8 Acknowledgements

In relation to the RNA sequencing of small extracellular vesicles, the authors gratefully acknowledge the University of Glasgow Shared Research Facility for their support and assistance with NGS library preparation, sequencing and preliminary analyses. Variant data for transcription factor analysis was obtained from the Rat Genome Database.

## 11 Supplemental Material

Supplemental methods Figures S1-4

